# Histone H3 tail modifications regulate structure and dynamics of the H1 C-terminal domain within nucleosomes

**DOI:** 10.1101/2023.05.11.540398

**Authors:** Subhra Kanti Das, Ashok Kumar, Fanfan Hao, Amber R. Cutter DiPiazza, Tae-Hee Lee, Jeffrey J. Hayes

## Abstract

Despite their importance, how linker histone H1s interact in chromatin and especially how the highly positively charged and intrinsically disordered H1 C-terminal domain (CTD) binds and stabilizes nucleosomes and higher-order chromatin structures remains unclear. Using single-molecule FRET we found that about half of the H1 CTDs in H1-nucleosome complexes exhibit well-defined FRET values indicative of distinct, static conformations, while the remainder of the population exhibits dynamically changing values, similar to that observed for H1 in the absence of nucleosomes. We also find that the first 30 residues of the CTD participate in relatively localized interactions with the first ∼20 bp of linker DNA, and that two separate regions in the CTD contribute to H1-dependent organization of linker DNA, consistent with some non-random CTD-linker DNA interactions. Finally, our data show that acetylation mimetics within the histone H3 tail induce decondensation and enhanced dynamics of the nucleosome-bound H1 CTD. (148 words)

## Introduction

In eukaryotes, chromatin condenses genomic DNA within the nucleus and regulates DNA accessibility through a variety of mechanisms to control gene expression. The primary structure of chromatin is formed by organizing sequential ∼200 bp segments of genomic DNA into nucleosomes, wherein ∼147 bp of DNA is wrapped about 1 ¾ times around an octamer of core histone proteins to form nucleosome cores, that are joined by a linker DNA segment of variable length (Cutter and Hayes, 2015; van Holde, 1989). In metazoans most nucleosomes are bound by a single molecule of a linker histone (H1), which stabilizes the nucleosome and also promotes the formation of the condensed chromatin structures that comprise chromosomes (Cutter and Hayes, 2015). Consequently, H1s are essential proteins that play critical roles in diverse nuclear processes including the formation of pericentric heterochromatin, silencing of transposable elements, maintaining the epigenetic landscape, and gene regulation (Kavi et al., 2016; Lu et al., 2013; Pan and Fan, 2016; Willcockson et al., 2021).

In higher eukaryotes, H1s have a tripartite structure composed of a central ∼80 residue trypsin-resistant central globular domain (GD) bounded by a short (20-35 residue) N-terminal domain (NTD), and a longer ∼100 amino acid residue C-terminal domain (CTD) (Cutter and Hayes, 2015) (Fig. 1). The central GD is more highly conserved among H1 subtypes than either the N- or C-terminal domains and adopts a ‘winged’ helix-turn-helix DNA binding motif (Cerf et al., 1993; Ramakrishnan et al., 1993) that interacts with the open minor groove at the nucleosome dyad, and with the two linker DNAs entering and exiting the nucleosome (Bednar et al., 2017; Zhou et al., 2015). The globular domain is responsible for structure-specific recognition and binding to nucleosome (Allan et al., 1986). Despite several high-resolution structures of H1-nucleosome complexes, the structure and interactions of the H1 NTD and CTD within nucleosomes remain poorly understood (Bednar *et al*., 2017; Hao et al., 2021; Zhou *et al*., 2015). While the NTD is unstructured in solution, it has been predicted to adopt some secondary structures when bound to the nucleosomes (Sridhar et al., 2020; Vila et al., 2001). Although the primary sequences of H1 CTDs are not well conserved, amino acid compositions are strikingly similar across species and subtypes, with about 40% of residues represented by basic amino acid residues, primarily lysines (Happel and Doenecke, 2009), consistent with the role of the H1 CTD in neutralizing the charge of DNA within condensed higher order chromatin structures (Allan *et al*., 1986; Bates et al., 1981; Clark and Kimura, 1990). Of note, H1 CTDs have sequence content emblematic of the class of intrinsically disordered proteins (Hansen et al., 2006; Ward et al., 2004), and are unstructured when free in aqueous solution (Bradbury et al., 1975). Moreover, peptides derived from the H1 CTD exhibit canonical secondary structures in structure-stabilizing solvents, or upon interaction with DNA or nucleosomes (Caterino et al., 2011; Clark et al., 1988; Fang et al., 2012; Roque et al., 2005; Zhou et al., 2021). These results are consistent with the idea that H1 CTDs are intrinsically disordered domains that acquire defined structure(s) upon interaction with target macromolecules (Bondos et al., 2022; Hansen *et al*., 2006). The disorder-to-order transition might be functionally important for chromatin structure in providing a large extent of charge neutralization but while tempering H1 binding affinities.

**Fig. 1:**
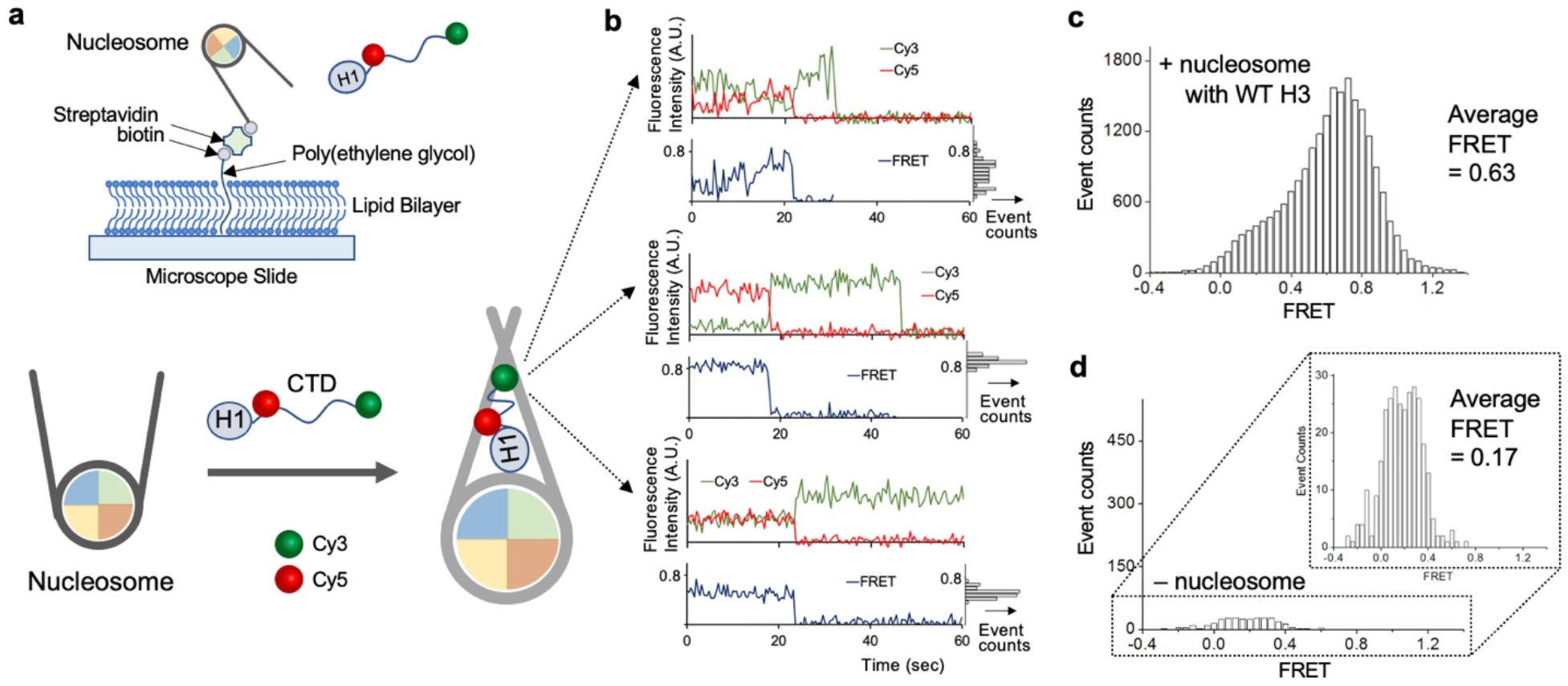
Single-molecule FRET (smFRET) reveals an ensemble of many related but distinct and rapidly interconverting condensed states of nucleosome-bound linker histone H1.0. **a** Experimental setup to investigate the conformations and dynamics of linker histone H1.0 CTD bound to a nucleosome. The CTD of H1.0 is labeled with the FRET pair Cy3 and Cy5 at the 101^st^ and 195^th^ amino acid residues. Nucleosomes are immobilized on a microscope slide coated with a lipid bilayer and FRET signals are collected from the slide surface via excitation with a total internally reflected laser beam at 532 nm. **b** Examples of fluorophore intensity and smFRET time traces from H1.0 CTD bound to the surface-immobilized nucleosomes. Fluorescence intensities are plotted as arbitrary units (A.U.). The FRET signals from an ensemble of H1.0 molecules are parsed into dynamic (top) or static (center and bottom) populations. **c** A histogram of FRET efficiencies from an ensemble of 335 nucleosome-bound H1.0 molecules reveals a wide distribution of CTD conformations with dominant higher FRET populations. The histogram is constructed by combining both the dynamic and static populations **d** A minimal number of H1.0 molecules associate with the surface in the absence of nucleosomes, and exhibit low FRET values. The total observation surface area was kept constant between **c** and **d**.

Indeed, it has been reported that the H1 CTD undergoes a substantial condensation upon canonical H1 binding to nucleosomes, consistent with a disorder-to-order transition resulting in a structured state or ensemble of structured states (Caterino *et al*., 2011; Fang *et al*., 2012; Fang et al., 2016; Hao *et al*., 2021). In addition, peptides derived from various H1 CTDs have been shown to adopt secondary structures upon interaction with DNA, or in structure-stabilizing solvents (Clark *et al*., 1988; Roque *et al*., 2005; Roque et al., 2012). On the other hand, investigations of H1 in complex with prothymosin-α (ProTα) or nucleosomes indicate that despite high affinity binding, the H1 CTD remains highly dynamic, which may facilitate rapid association/dissociation of H1 by chaperones (Borgia et al., 2018; Heidarsson et al., 2022; Sottini et al., 2020), while NMR investigations of an H1 CTD peptide in complex with DNA indicate that the CTD remains disordered within the complex (Turner et al., 2018). Moreover, epigenetic modifications within the H3 tail domain alter overall H1 CTD condensation but it is unclear whether these modifications influence H1 CTD dynamics. To investigate the extent of condensation of the nucleosome-bound H1 CTD and the effect of H3 tail modifications, we performed single-molecule Förster Resonance Energy Transfer (FRET) studies and examined the distribution and interconversion of structural states.

We find that the nucleosome-bound H1 partitions between stably condensed states exhibiting well-defined FRET values, and dynamic states displaying interconversion between condensed conformations. Modifications of H3 tail domain skew the H1 to more dynamic and less condensed FRET states. In addition, crosslinking assays and assessments of linker DNA reorganization indicate that to some extent specific regions within the H1 CTD participate in localized interactions with the linker DNA.

## Results

The intrinsically disordered linker histone H1.0 CTD condenses upon binding to nucleosomes, perhaps indicative of adoption of a structure or an ensemble of structures in the nucleosome environment (Fang *et al*., 2012; Fang *et al*., 2016; Heidarsson *et al*., 2022). To better understand the structural state(s) adopted by the nucleosome-bound H1.0 CTD, we employed single-molecule FRET (smFRET). Nucleosomes were reconstituted with a 207 bp biotinylated DNA fragment, and incubated with Cy3 and Cy5 labeled H1.0 G101C K195C, wherein the fluorophores were attached to residues bracketing the CTD domain (Fig. 1a). The H1-bound nucleosomes were then injected into a flow channel and allowed to bind to a streptavidin-coated surface (see Methods). Unbound nucleosomes were flushed with buffer, and the surface was illuminated with a 532 nm laser and the FRET signals from individual H1.0 molecules were recorded.

Numerous fluorescence spots exhibiting FRET were identified in each experiment, and Cy3 and Cy5 emission from individual spots was recorded for 3-5 minutes and a histogram of FRET responses compiled (e.g. Fig. 1, b and c). We found that H1.0 with Cy3 and Cy5 labels at each end of the CTD exhibited an average overall FRET efficiency of 0.63, in the presence of nucleosomes (Fig. 1c), consistent with earlier ensemble FRET measurements (Fang *et al*., 2012; Fang *et al*., 2016; Hao et al., 2020). In the absence of nucleosomes, a much lower number of H1s non-specifically associated with the passivated surface, as indicated by the total number of event counts (Fig. 1d). This population exhibited an average FRET efficiency of 0.17, again in agreement with ensemble FRET measurements for free H1 and indicating that the CTD of H1 non-specifically associated with the passivated surface exhibits FRET consistent with a random coil conformation (Fang *et al*., 2012; Fang *et al*., 2016). Moreover, a comparison of the total event counts for nucleosome-bound and unbound H1 indicates that only about 5.5% to the nucleosome-bound H1 signal is contributed by H1 non-specifically bound to the passivated surface (Fig. 1, c and d). In sum, these data are in agreement with ensemble and smFRET experiments indicating that the H1 CTD undergoes significant condensation upon H1 binding to nucleosomes (Caterino *et al*., 2011; Fang *et al*., 2016; Heidarsson *et al*., 2022).

As is clear in Fig. 1c, the FRET distribution from H1-nucleosome complexes does not follow a standard function representing a single structural state whose statistics are well defined (e.g., a single gamma or gaussian distribution). However, this distribution of FRET efficiencies serves as a visual guide representing the distribution of CTD conformations within nucleosome-bound H1s. The distribution could represent an assemblage of rapidly interconverting structures or static structures, or some combination of both. Inspection of the smFRET traces we obtained for nucleosome-bound H1 shows that about half of the traces exhibit only one FRET state, typically lasting for 10’s of seconds, before photobleaching, while the remaining traces clearly exhibit significant dynamics, with multiple FRET values in the traces (Fig. S1). We classify the former as “static” and the latter as “dynamic” and plotted a histogram of each group independently (Fig. 2 b-c). Interestingly, the static (non-interconverting) population exhibited an average FRET of 0.73, while the dynamic population exhibited an average FRET of 0.55, and contains greater counts of low-FRET values (Figs. 2 b-c). To better understand the relationship between interconverting states in the dynamic traces, we plotted transitions between the FRET states (Fig. S2). The scatter plot shows no clustering of FRET values or defined relationships between initial and final FRET values, indicating the dynamic population is comprised of complexes randomly interconverting between a wide range of FRET states. Similarly, dynamic behavior was found for nearly all (>95%) H1s non-specifically associated in the absence of nucleosomes (see Fig. 1d) as well as for antibody-tethered FLAG-tagged H1 (Fig. S3).

**Fig. 2:**
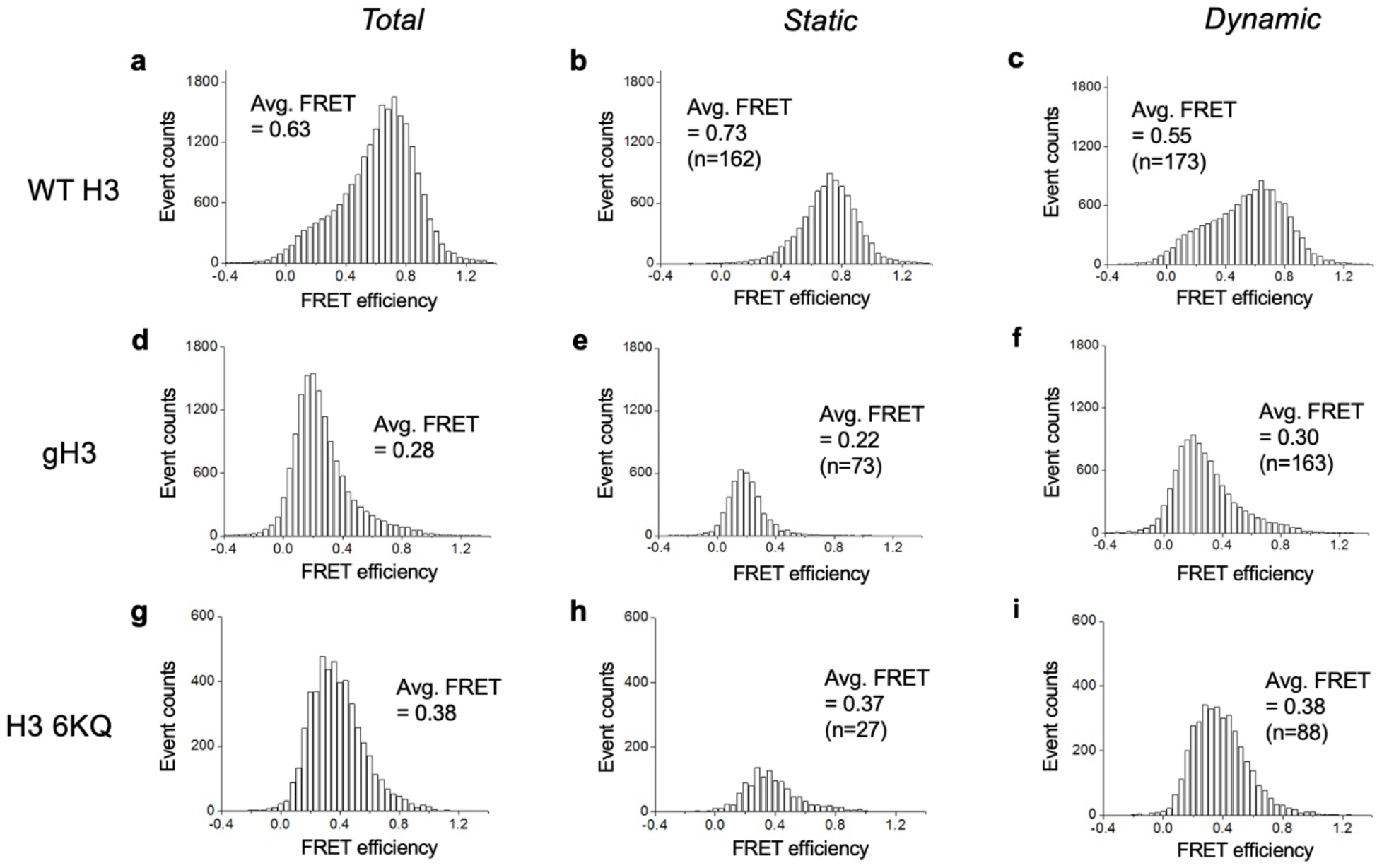
Modifications within the H3 tail domain alter the extent of condensation and dynamics of nucleosome-bound H1 CTD. FRET efficiency histograms representing all nucleosomes (a, d, g), the static population (b, e, h) and the dynamic population (c, f, i) for H1.0 bound to wildtype H3 (WT H3, a-c), nucleosomes lacking H3 N-tail domains (gH3, d-f), and nucleosomes with 6 lysine residues in the H3 N-terminal tail substituted by glutamine as mimetics of lysine acetylation (H3 6KQ, g-i) are shown. The number of H1.0 molecules observed (n) is indicated. Note data shown in Fig. 1c is repeated in 2a for reference.

To better understand the H1 conformations that comprise the static population, we examined their FRET traces. A set of traces with average FRET values spanning the total distribution are shown in Fig. S4. The average FRET efficiencies and the standard deviations of the FRET distributions of the individual molecules from the entire population of static H1 CTDs are shown in Fig. S5. This analysis shows that the aggregate FRET response (Fig. 2b) is a convolution of individual H1-nucleosome complexes exhibiting narrow FRET distributions with peaks clustering mostly in the high FRET range. The narrow FRET distributions of the vast majority indicate that the static population is comprised mainly of distinct and stable FRET structures. Overall, this data shows that about half of the of nucleosome-bound H1 CTDs are represented by a family of stable, condensed high-FRET species.

We previously discovered that removal of the histone H3 tail domain, but not other core histone N-tail domains, significantly reduces the overall condensation of the H1 CTD (Hao *et al*., 2020).

Moreover, we find that mimics of epigenetic acetylation within the H3 tail similarly affect H1 CTD structure. However, the effects on H1 CTD structure do not appear to be due to changes in linker DNA trajectory in the H1-bound nucleosome (Hao *et al*., 2020). To gain a better understanding of the mechanism by which H3 tail modifications affect the condensed state of the H1 CTD, we repeated the smFRET analysis with nucleosomes containing either unmodified H3 (WT H3), tailless H3 (gH3), or H3 6KQ, in which glutamine was substituted as a mimetic for six lysines known to be acetylated in the H3 tail domain *in vivo*. The nucleosomes were incubated with H1 in the same manner as the native nucleosomes and smFRET profiles determined. We find, in accordance with prior work (Hao *et al*., 2020), that removal of the H3 tail domain or installation of acetylation mimetics significantly reduces the extent of H1 CTD condensation upon binding to nucleosomes, with nearly no H1 CTDs populating the high FRET states adopted by H1 CTDs bound to unmodified nucleosomes (Fig. 2, d and g, Figs. S4 and S5). Averages of FRET responses for the population of H1-nucleosome complexes shift from 0.63 for nucleosomes containing unmodified H3 (Fig. 1) to 0.28 or 0.38 for nucleosomes containing tailless H3 or H3 K6Q nucleosomes, respectively (Fig. 2, a, d, and g). Thus, modifications of the H3 tail domain induce a much less condensed H1 CTD compared to that found in the WT nucleosome-H1 complex. Moreover, we observed that the fraction of H1 CTDs exhibiting dynamic interconversion between FRET states rises substantially upon H3 tail modification, from 52 ± 3% for nucleosomes containing WT H3, to 69 ± 3% and 77 ± 5% for gH3 and H3 K6Q nucleosomes, respectively (Fig. 2 and Table 1).

**Table 1:**
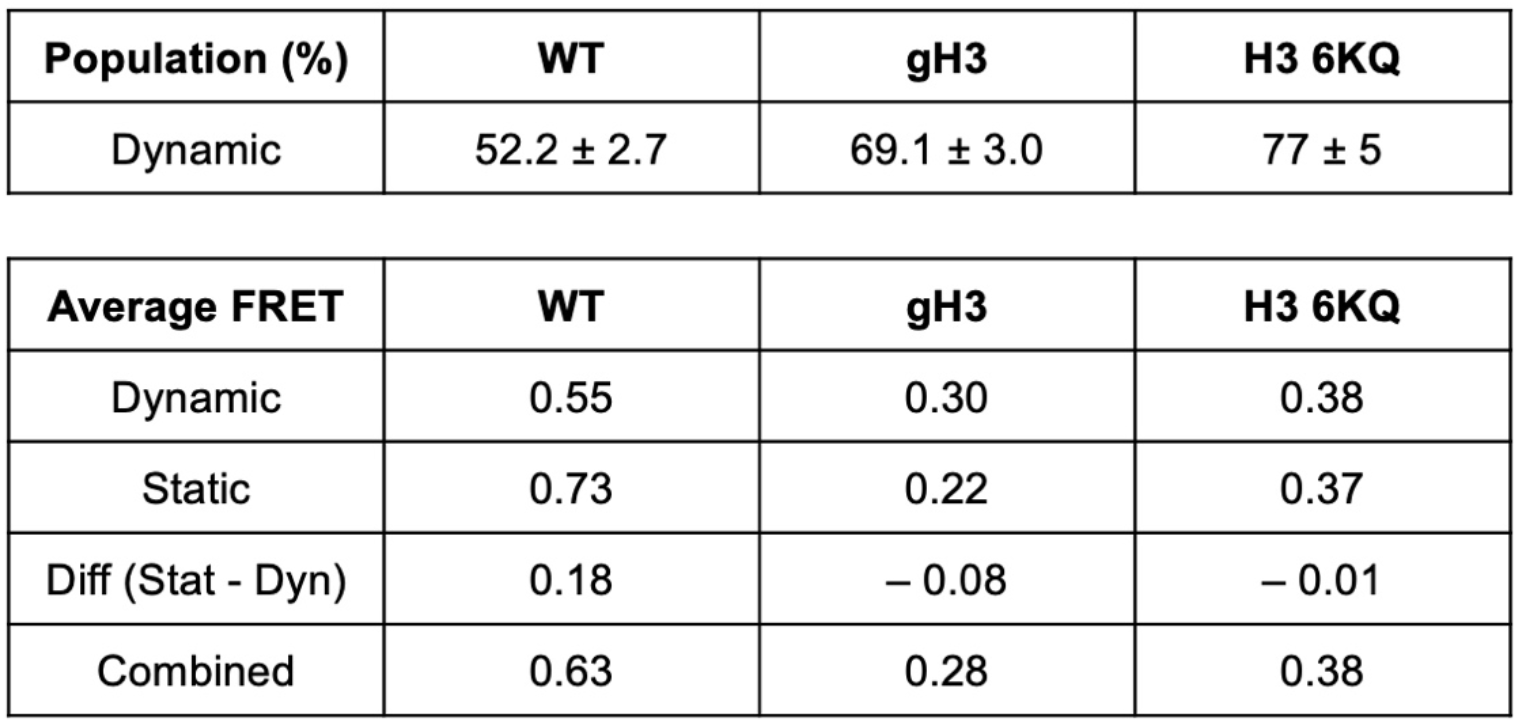
The population densities and average FRET efficiencies of dynamic and static H1-CTD populations in WT, gH3, and H3 6KQ nucleosomes. The errors in the population densities are the binomial distribution errors from the total sample sizes of 335, 236, and 115 in the WT, gH3, and H3 6KQ cases.

| **Population (%)** | **WT** | **gH3** | **H3 6KQ** |
| --- | --- | --- | --- |
| Dynamic | 52.2 ± 2.7 | 69.1 ± 3.0 | 77 ± 5 |

| **Average FRET** | **WT** | **gH3** | **H3 6KQ** |
| --- | --- | --- | --- |
| Dynamic | 0.55 | 0.30 | 0.38 |
| Static | 0.73 | 0.22 | 0.37 |
| Diff (Stat - Dyn) | 0.18 | – 0.08 | – 0.01 |
| Combined | 0.63 | 0.28 | 0.38 |

The smFRET results indicate that the nucleosome-bound H1 CTD adopts a broad family of distinct condensed conformations. As such, CTD interactions with linker DNA may be essentially diffuse or, alternatively, some portion of this domain may adopt more localized contacts. To probe the potential interactions between the H1 CTD and linker DNA we performed site-directed crosslinking. Cysteine residues substituted at positions 101, 112, 130 and 173, were labeled with a UV-activatable crosslinking probe azidophenacyl bromide (APB, Fig. 3a), and crosslinks to DNA within the H1-nucleosome complexes mapped to single-nucleotide resolution. Both WT H1 and APB-labeled mutants showed stoichiometric binding to nucleosomes (Fig. 3b, top). Moreover, the APB-modified mutants, but not the unmodified WT H1 efficiently formed covalent crosslinks to nucleosome DNA upon UV irradiation (Fig. 3b, bottom), consistent with previous work (Bednar *et al*., 2017). We mapped the location of the crosslinks on denaturing gels, and found bands indicating localized crosslinking between G101C, I112C and S130C at positions 81, 83/84, and 94/96 base pairs from the nucleosome dyad, respectively (Figs. 3c and S6).

**Fig. 3:**
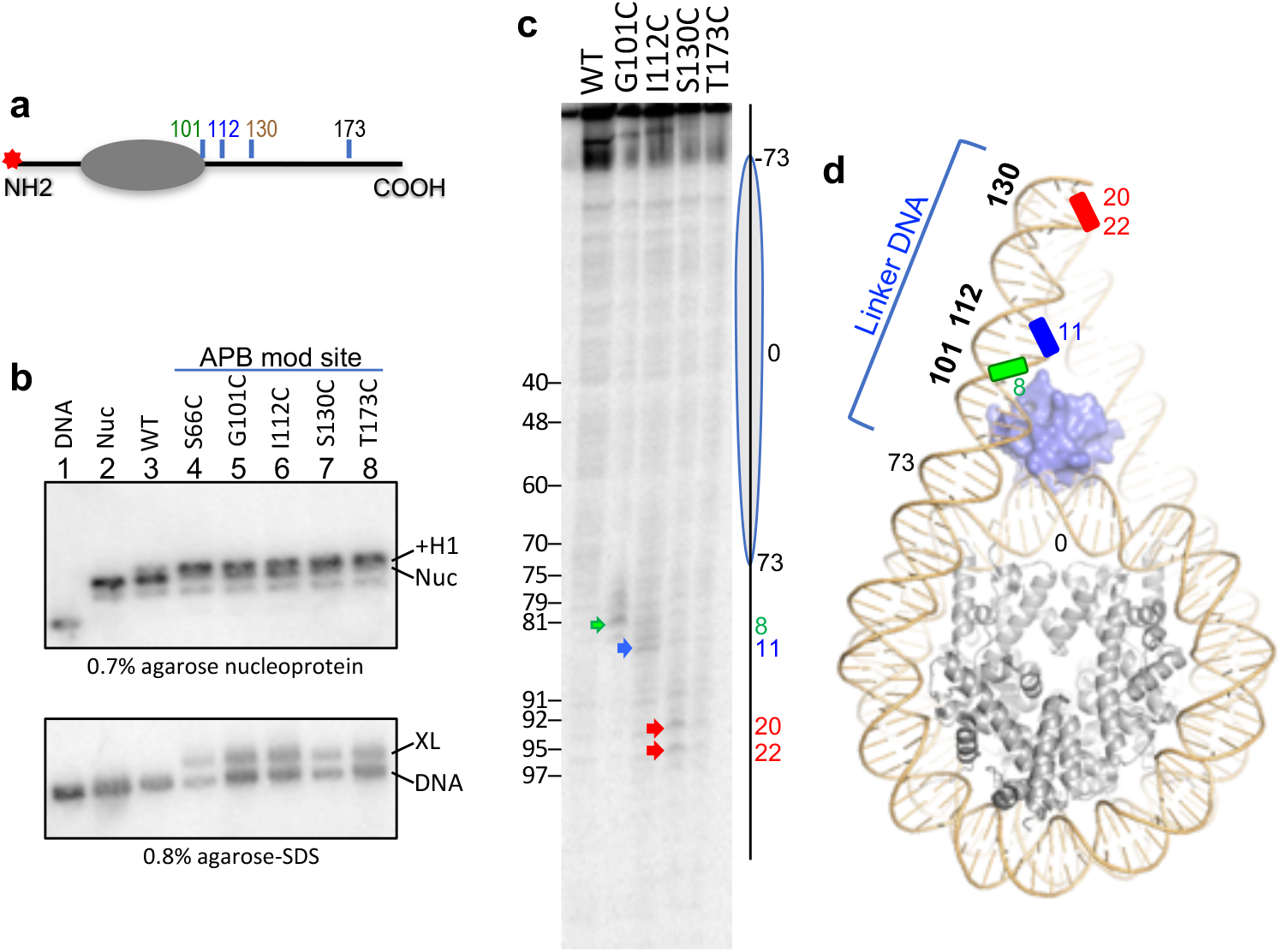
Crosslink mapping of sites of association between the H1 CTD and linker DNA. **a** Positions of cysteine substitutions modified with APB in the indicated H1 mutants. The position of the radiolabel on the 5’ end of the top strand is indicated by the red star. **b** *Top* H1 WT and APB-modified mutants bind 601 nucleosomes. The band corresponding to the nucleosome (Nuc) and H1-nucleosome complex (+H1) are indicated. *Bottom* APB-modified H1s crosslink to nucleosome DNA. Bands corresponding to free DNA (DNA) and DNA crosslinked to modified H1s (XL) are indicated. **c** Example of crosslink mapping on nucleosome DNA. Sites of crosslinking for G101C-APB, I112C-APB, S130C-APB are indicated by the green, blue and red arrows, respectively. No discrete crosslinking is detected for T173C-APB. The nucleosome core (oval) and linker DNA (line) are indicated as are positions in the 601 DNA, indicated as distance from the nucleosome dyad position (Bednar *et al*., 2017). **d** Schematic showing sites of crosslinking in the nucleosome DNA (PDB ID: 5NL0).

However, no significant crosslinking sites were able to be identified for H1 T173C-APB, despite robust crosslinking to nucleosome DNA (Figs. 3b and c). We note that previous work shows that H1.0 binds to the 601 nucleosome in both ‘on-dyad’ orientations in the canonical nucleosome binding pocket, with residues 66 and 101 making symmetrical interactions with the linker DNA (Bednar *et al*., 2017). Plotting the position of crosslinked DNA sites within a model of an H1.0-nucleosome structure (Bednar *et al*., 2017) indicates that the first ∼30 residues of the H1 CTD appear to track somewhat linearly along the first ∼20 bp of linker DNA.

To further define how the H1 CTD interacts with linker DNA, we examined the effect of truncations within the H1 CTD on linker DNA orientation within the nucleosome. Linker DNA within H1-lacking nucleosomes on average exist in an open conformation, but upon H1 binding are drawn together such that their end-to-end distance (E-T-E) is significantly reduced (Bednar *et al*., 2017). We measured the relative distance between linker DNA ends in the absence or presence of WT H1 and H1 CTD truncation mutants by ensemble FRET (Fig. 4) with Cy3 and Cy5 attached near the ends of the DNA fragment used for nucleosome reconstitution. As in previous work nucleosomes in the absence of H1 exhibited a low level of FRET due to the divergent angles of the linker DNAs, but FRET was significantly elevated upon the addition of WT H1, indicating much closer apposition of the two linker DNA ends (Fig. 4c). H1s with deletions of 22 (Δ 22), 44 (Δ 44), or 66 (Δ66) amino acid residues from the C-terminus of H1 yielded FRET responses significantly lower than that of WT H1, indicating increased distance between the ends of the linker DNAs. Further deletion of 88 (Δ88) residues, or the entire 96 residue CTD (NG) resulted in an additional decrease in FRET efficiency, to a level not significantly different from that of the nucleosome in the absence of H1s (Nuc) (Fig. 4c). These results suggest that two distinct segments of the H1 CTD, lying between residues 175 and the C-terminus of the protein, and between residues 109 and 130 play key roles in orienting the linker DNA.

**Fig. 4:**
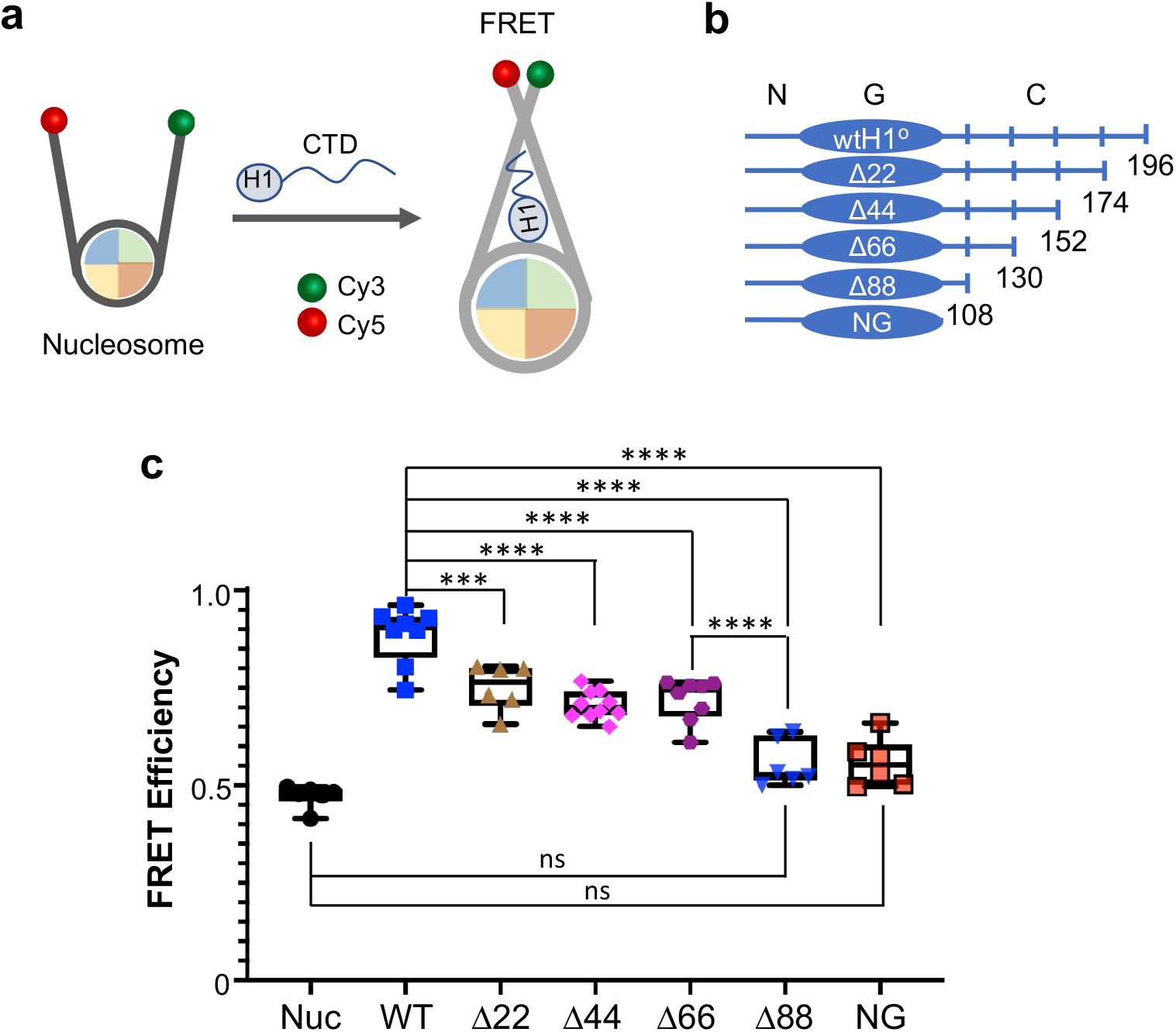
Two regions within the H1 CTD contribute to linker DNA organization within the nucleosome. **a** Experimental setup to monitor FRET from the linker DNA ends of nucleosomes. **b** Serial deletions within the H1 CTD, the N, G, and C are N-terminal domain, globular domain, and C-terminal domain, respectively. Numbers below indicate terminal residue in each protein. Red bars indicate regions that alter linker end-to-end distance upon deletion shown in c. **c** Plot of FRET efficiencies for nucleosomes bound by WT H1 and deletion mutants compared to the H1-free nucleosome (Nuc). NG indicates deletion of the entire 96-residue CTD, leaving only the N and G domains. N ≥ 6 for each determination. **** and *** indicate p-values of < 0.0001 and <0.001, respectively.

## Discussion

We find that the intrinsically disordered H1 CTD undergoes a drastic condensation upon binding to nucleosomes, with the majority of the population adopting high-FRET conformations. Moreover, the CTDs in approximately half of the H1-bound nucleosomes exhibit a highly dynamic character, in which FRET values interconvert on a time scale clearly observable at a 100 ms time resolution, while about half of the H1 CTDs exhibit markedly distinct FRET traces indictive of structures that are well-defined and stable over timescales of a few to 10s of seconds. Crosslinking studies show that the first ∼30 residues of the H1 CTD make relatively localized contacts with the first ∼20 bp of the nucleosome-proximal linker DNA, while the C-terminal region of the CTD likely interacts with multiple nucleosome DNA sites. Moreover, we find that two separate regions within the CTD contribute to linker DNA organization in the H1-bound nucleosome, consistent with the idea that regions within the CTD perform distinct functions in the formation of chromatin higher-order structures (Lu et al., 2009; Lu and Hansen, 2004).

The broad range of smFRET values observed for the H1–nucleosome complexes is consistent with a large number of conformations contributing to the ensemble of condensed H1 CTD structures. Analyses of the FRET histograms associated with static complexes reveal that the overall distribution is a convolution of distinct and stable FRET species with narrow FRET distributions, indicating that the static population contains many distinct structures. Interestingly, the static population is clearly biased toward high-FRET values, while the dynamic population shows a much greater tendency to form low-FRET complexes.

Our data show that inclusion of tailless H3 or H3 containing mimetics of epigenetic acetylation in nucleosomes strongly biases CTD conformations toward the low-FRET structures. Indeed, only H1 CTDs bound to WT H3 nucleosomes exhibit an appreciable amount of high FRET values. Moreover, the fraction of H1 CTDs existing in dynamic states increases drastically upon alteration of the H3 tail domain, from ∼50 % for WT nucleosomes to 70 and 80 % of the population for gH3 and H3 6KQ nucleosomes, respectively (Table 1). These results suggest that modifications in the H3 tail domain do not fundamentally alter the structural states explored by the H1 CTD but rather bias the distribution to more dynamic, extended conformations represented by the low-FRET states, potentially further increasing accessibility of the CTD to posttranslational modifiers and chaperones (Heidarsson *et al*., 2022). Indeed, previous work has shown that binding of the chromatin architectural transcription factor HMGN2, and phosphorylation mimics within the H1.0 CTD also influence H1 CTD structure in the nucleosome (Hao *et al*., 2021; Hao et al., 2022; Murphy et al., 2017). It will be interesting to determine the extent to which H1 CTD dynamics are altered by such epigenetic signals.

The H3 tail modifications employed in this work increase DNA unwrapping (Hao *et al*., 2020). Moreover, unwrapping enforced by binding of a transcription factor abrogates interaction with between the H1 CTD and the unwrapped linker, but does not result in loss of binding of the H1 globular domain at the nucleosome dyad (Burge et al., 2022), consistent with the asymmetric interaction of the globular domain with linker DNA (Bednar *et al*., 2017). These results suggest a model wherein the CTD interacts preferentially with a single (unwrapped) linker DNA, resulting in a less condensed and more dynamic ensemble of H1 CTD structures. Although the FRET data indicate that the condensed H1 CTD structure adopts a large collection of conformations exhibiting a wide range of FRET values, our results suggest that two distinct portions of the CTD play roles in orienting nucleosome linker DNA in nucleosomes. The effect of CTD deletions on linker DNA end-to-end distance indicate that the terminal ∼22 residues of the H1 CTD (residues 175 – 196) and residues 109-130 contribute each independently contribute to drawing the linker DNAs together. These results are consistent with previous reports indicating that a short region in the extreme N-terminal region of the CTD of human H1.5 and human H1.0a are essential for bringing the two linker DNAs together and forming the initial linker stem structure (Burge *et al*., 2022; Syed et al., 2010). A similar highly basic region exists within residues 109-130 of *Xenopus* H1.0a and likely is responsible for the effect observed here. Moreover, Lu et al. showed two discontinuous subdomains of H1 CTD are required for stabilizing the folded secondary chromatin structures and promoting the self-association of nucleosomal arrays into oligomeric tertiary chromatin structures, respectively (Lu *et al*., 2009; Lu and Hansen, 2004). In summary, our finding that two distinct regions within the H1 CTD contribute to linker DNA apposition does not support a simple, completely unstructured model for the H1 CTD, which predicts a graded loss of function with incremental truncation of the H1 CTD.

In addition, site-directed crosslinking experiments via selected sites within the H1 CTD detect localized interactions for three sites spanning the first 30 residues at the N-terminal region of the CTD domain, suggesting some discrete structure exists in this region. However, no specific or localized crosslinking was detected for H1 T173C-APB, despite a comparable yield of crosslinked species, suggesting that these crosslinks are spread across a wider range of positions in the nucleosome DNA. Thus, our data suggest that at least the first ∼30 residues of this domain appear to be at least partially localized with respect to linker DNA. These results are consistent with a recent multi-pair H1-nucleosome DNA FRET approach that indicated the N-terminal portion of the CTD is mostly juxtaposed to a labeled position about 14 residues beyond the edge of the nucleosome core region (Heidarsson *et al*., 2022). Moreover, FRET studies show that the H1 CTD exists as an extended random coil in aqueous solution, with a calculated distance between residues 101 and 195 is ∼8 nm while a FRET pair installed at residues 101 and 173 are separated by a shorter average distance of ∼6.2 nm. However, upon association with nucleosomes the distance between residue 101 and 173 is ∼4.9 nm, longer than that between G101C and K195C (∼4.2 nm) (Caterino *et al*., 2011; Fang *et al*., 2012). These results provide support for the idea that the H1 CTD exhibits some level of structural order in the nucleosome-bound state, consistent with a recent model derived from the multiplexed FRET pair study (Heidarsson *et al*., 2022). The electrostatic neutralization of the ∼40 excess positive charges within on the H1 CTD upon interaction with the linker DNA might elevate the hydrophobicity of the H1 CTD overall and facilitate its folding although general charge shielding alone from an ionic environment is not sufficient to induce specific H1 CTD folding (Fang *et al*., 2016). On the other hand, the nucleosome-induced condensation of the H1 CTD possibly allows attenuation of the binding affinity of H1 to the nucleosome, allowing rapid equilibration about the nucleus, on a minutes timescale. Second, the dynamic nature of the disordered region can serve as a platform for regulation by posttranslational modifications (PTMs) (Borgia *et al*., 2018; Hansen *et al*., 2006; Sottini *et al*., 2020), allowing ready access to trans-acting factors, including DNA methyltransferases DNMT3B and DNMT1 (Yang and Hayes, 2011) and the chromatin architectural factors HMGN1/2 (Murphy *et al*., 2017).

We define the structure of the nucleosome-bound H1 CTD as an ensemble of distinct structures with FRET distributions biased toward a high-FRET value indicating overall condensation of this domain which become less condensed and more dynamic upon H3 tail acetylation. Nevertheless, our data indicate that a ∼30 residue section of the CTD closest to the globular domain exhibits localized interactions with the nucleosome proximal linker DNA, and initiates linker DNA reorganization. In prior work, we showed that linker DNA trajectory in turn can influence H1 CTD structure and changes in linker DNA trajectory associated with early states of folding alter the final condensed state of the H1 CTD. Therefore, it will be of interest to define the distributions of states and dynamics of H1 CTD bound to oligonucleosome arrays during chromatin folding.

## Acknowledgements

The work was supported by National Institutes of Health Grants R01GM052426 and R35GM149420 (to J.J.H), T32GM068411 (to A.R.C.D.), R01GM123164 and R01GM130793 (to T.L). We also acknowledge support from NIH S10-OD021489-01A1.

## Data Availability Statement

All data will be made available upon request.

## Methods

### Protein preparation and nucleosome reconstitution

Linker histone *Xenopus laevis* H1.0 (hereafter referred to as H1) and H1 G101C K195C, in which cysteines are located at both ends of the H1 CTD, as well as core histones were expressed in bacterial cells and purified as described (Hao *et al*., 2020). The FLAG-tagged version of WT H1 (FLAG-H1) G101C K195C was generated using a Qiagen Q5 site-directed mutagenesis kit with FP 5’ATCTGGACGGTGCAAGTAAGGATCCC3’ and RP 5’TTCTTTGGGCTGGCTTTTG’ and the pET3aWTH1.0 G101C K195C vector (Caterino *et al*., 2011), verified by DNA sequencing. The 207 bp DNA fragments containing the 601 nucleosome positioning sequence used for nucleosome reconstitution were isolated by digestion of plasmid p207-12 with EcoRV as described (Hao *et al*., 2020). Nucleosomes were reconstituted via a standard salt dialysis. Briefly, 5 μg of H3/H4 tetramer, 5.5 μg H2A/H2B dimer and 10 μg 601 DNA were mixed in reconstitution buffer (10 mM Tris, pH 8.0, 1 mM EDTA, 5 mM DTT and 2 M NaCl) in a total volume of 300 μL, transferred to a dialysis tubing, then dialyzed against buffers containing 10 mM Tris-HCl, pH 8.0, 1 mM EDTA, and decreasing concentrations of NaCl (1.2 M, 1 M, 0.8 M, 0.6 M), for 2 hrs at 4°C, followed by dialysis against TE overnight at 4°C. Reconstituted nucleosomes were purified on 5 ml 7%-20% sucrose/TE gradients by ultracentrifugation in a Beckmann SW41 rotor for 18 h at 34,000 x g at 4°C. The fractions containing nucleosomes were combined and concentrated using a microfuge tube filtration unit with a Nominal Molecular Weight Limit of 50 kDa (EMD Millipore).

Nucleosome fractions were analyzed on 0.7% native agarose gels.

### Cy3 and Cy5 labeling

H1 G101C K195C was incubated in 50 mM DTT for 1 h on ice, then DTT was removed by Bio-Rex chromatography and fractions immediately frozen on dry ice (Hao *et al*., 2020). Reduced H1 G101C K195C was treated with ∼5-fold molar excess of either maleimido-Cy3, or maleimido-Cy5, or a 50/50 mix of both (GE Healthcare, PA23031 and PA25031) for 30 min at room temperature in the dark. Free dyes were removed by another round of Bio-Rex chromatography. Concentration of fluorophore labeled H1 was determined by quantitative comparison with an H1 standard on SDS-PAGE gels and the efficiency of labeling was determined by comparison to total fluorescence of dye standards and known labeling controls and is typically ∼90% (Hao *et al*., 2020).

### Microscope slide preparation

We coated the surface of a quartz microscope slide with a sub-monolayer of biotin-PEG-silane followed by lipid bilayer deposition according to previously published protocols (Yue et al., 2016). Briefly, a sub-monolayer of biotin-PEG-silane (MW 3400, Laysan Bio, Arab AL) was grafted on a scrupulously cleaned quartz microscope slide. For lipid bilayer deposition, lipid vesicles extruded through an 80 nm pore-sized membrane filter (Avanti Polar Lipids, Alabaster AL) were prepared from 1,2-Dioleoyl-sn-glycero-3-phosphoethanolamine-N-[methoxy(polyethylene glycol)-5000] (ammonium salt) (DOPC) (Avanti polar lipids, Alabaster AL). Flow channels were constructed on the surface of a biotin-PEG coated slide according to previously published protocols (Lee et al., 2019). A flow channel was filled with the vesicle solution and incubated for 45 minutes to deposit a lipid bilayer.

### Single-molecule measurements

A flow channel was incubated with 0.1 mg/mL streptavidin (Sigma-Aldrich) solution for 2 minutes for streptavidin-biotin conjugation. The flow channel was then washed with TE50S buffer (10 mM Tris (pH 8.0), 1 mM EDTA, and 50 mM NaCl) to flush the unbound streptavidin molecules. The nucleosomes (WT, gH3, or H3 6KQ) labeled with biotin at one end of the DNA were incubated with Cy3/Cy5 labeled H1 at a molar ratio of 1:5 (2 nM nucleosome and 10 nM H1) for 5 minutes in the TE50S buffer. The incubated sample was diluted to 100 pM nucleosome concentration and injected inside the flow channel to immobilize the nucleosome-H1 complexes via streptavidin-biotin conjugation. The flow channel was incubated for 2 minutes and then washed with 50uL of imaging buffer (10 mM Tris (pH 8.0), 1 mM EDTA, 50 mM NaCl, 2mM Trolox, 7mM protocatechuic acid (PCA), 0.7 mM protocatechuate-3,4-dioxygenase (PCD)) to eliminate all the surface unbound nucleosomes from the flow channel. The addition of Trolox, PCD, and PCA is to elongate the lifetimes of the fluorophores and reduce their blinking. Fluorescence signals from the samples were taken using a lab-built total internal reflection (TIR) fluorescence microscope based on a commercial microscope (Nikon, TE2000) in a movie format with a frame rate of 10 frames/second as previously published (Lee *et al*., 2019). Briefly, the surface of a flow channel was illuminated with a 532 nm Laser (150 mW, CrystaLaser) in a prism-coupled TIR geometry and the fluorescence images from the microscope were recorded with an EMCCD camera (IXON Ultra 897, Oxford Instruments) until most of the Cy3 fluorophores photobleached. Optical filters were used to filter the Cy3 signal (650DCXR and HQ550LP, Chroma Technology) and Cy5 signal (650DCXR and HQ650LP, Chroma Technology). Intensities of Cy5 and Cy3 are corrected for 4 % bleed-through from the Cy3 to the Cy5 channel. Direct excitation of Cy5 at 532 nm is negligible above the background noise level in our setup. The control single-molecule FRET (smFRET) measurements from non-specifically surface-bound H1 in the absence of any nucleosome were carried out under identical experimental conditions including the total observation time and surface area as those in the case with the wildtype H1 bound with nucleosomes. The control smFRET measurements from nucleosome-free FLAG-H1 immobilized on the surface via FLAG-anti-FLAG interactions were carried out under identical conditions with some minor modifications as follows. To immobilize the anti-FLAG antibody, we first deposited streptavidin on the slide as described above, and then, incubated 0.1 mg/mL biotinylated protein A (Thermo Fisher Scientific) in a flow channel for 15 min, which was followed by a wash with TBS buffer (50 mM Tris-HCl pH 7.6, 150 mM NaCl) and subsequent incubation with 300 nM anti-FLAG antibody (monoclonal ANTI-FLAG® M2 from Sigma) for 120 min. After washing the free anti-FLAG antibody with TBS buffer, FLAG-H1 at 100 pM was subsequently incubated in the flow channel for 4 min followed by another washing with the imaging buffer. In each flow channel, fluorescence movies of 3-5 min length were taken during the first 30 minutes after sample immobilization. On average, each movie shows 221 ± 20 fluorescent spots detected in the Cy3 channel out of which 112 ± 17 exhibit single-step photobleaching. The other spots show two or more photobleaching steps, indicating more than one Cy3, and therefore, were excluded from further analysis. Out of the single Cy3 spots, 83 ± 5 % show FRET signals, indicating a Cy3-Cy5 pair. The deviation from 100 % is likely due to contaminant fluorescent spots miscounted as Cy3 spots and premature photobleaching of Cy5. We also note that some of these FRET traces might include signals from nucleosomes bound with more than one H1 where only one is labeled with Cy3 and Cy5 while the others are labeled with two Cy5 molecules. However, since random labeling of the two sites that should result in 50 % Cy3/Cy5, 25 % Cy3/Cy3, and 25 % Cy5/Cy5 per H1, the maximum population density of these cases is only 8.6 ± 2.0 % of the single Cy3 spots based on our previously published ratio of ∼2:1:1 for one:two:three H1s bound per nucleosome (Yue *et al*., 2016). These cases should show multiple Cy5 photobleaching steps if the other Cy5 molecules form inter-molecular FRET. Therefore, we excluded any spot showing multiple photobleaching steps for Cy5 from further analysis. Finally, we also excluded FRET traces showing a signal-to-noise ratio lower than 4.0 at 100 ms integration. These filtering criteria resulted in selection of 48 ± 11 % of the single Cy3 spots for further analysis.

### Single-molecule data analysis

The first 5 frames of each fluorescence movie file were analyzed to identify Cy3-Cy5 fluorophore pairs from individual nucleosome-H1 complexes. From each complex, Cy3 and Cy5 fluorescence intensities were extracted for all the movie frames, which result in a time series of their intensities. With the result, a time trace of the FRET efficiencies was constructed using an approximate FRET efficiency formula of E = I_Cy5_ / (I_Cy3_ + I_Cy5_) at every time point, where E is the FRET efficiency and I_A_ is the intensity of fluorophore A. These FRET efficiencies were combined to construct a FRET efficiency histogram. Note that we did not seek accurate measurements of inter-fluorophore distances from FRET efficiencies. Instead, we measured the FRET efficiencies to make internal comparisons of inter-fluorophore distances among the cases of WT, gH3, and H3 6KQ nucleosomes.

### Linker DNA FRET analysis

The linker DNA end-to-end distance FRET experiments were performed as described (Hao *et al*., 2020). Briefly, FRET-labeled 207 bp DNA was prepared by PCR using labeled primers (GenScript) ATCGGACCC/iCy5N/ATACGCGGCC (forward primer), and AGTAG/iCy3N/ATTAATTAATATGAATTCGGATCCACATGCAC (reverse primer), in which Cy3 and Cy5 were conjugated to amino-modified C6-dT at internal positions to avoid fluorophore end-stacking.

Fluorescently labeled 207 bp DNA fragments were purified with a PCR purification kit (Qiagen), and nucleosome reconstitution performed as described above. Unlabeled H1.0 was titrated into labeled nucleosomes over a narrow range to ensure 1:1 binding, and ensemble FRET determined as described (Hao *et al*., 2020), with the fraction of donor labeled molecules d^+^ = 1, and extinction coefficients for the Cy3 donor: (515 nm) = 53160 cm^-1^M^-1^, and for Cy5 acceptor: (515) = 3749 cm^-1^M^-1^ and (610) = 118400 cm^-1^M^-1^. The data were analyzed and plotted using Prism-GraphPad software and means compared using Tukey’s multiple comparisons test; (***) P-value <0.001, (****) p-value <0.0001, (ns) p-value >0.1.

### H1-nucleosome DNA Crosslinking

H1 mutants containing an N-terminal FLAG tag and cysteines at positions indicated in Fig. 3 and S6 were generated and modified with azido-phenacylbromide (APB) as previously published (Bednar *et al*., 2017). Large scale, 500 μl H1 binding reactions containing ∼200,000 cpm (∼0.25 nmol) radiolabeled nucleosome were assembled, and 10μl samples analyzed on native nucleoprotein gels to ensure stoichiometric H1-nucleosome binding, and the remaining portions of the reactions were irradiated with UV light as described (Bednar *et al*., 2017). Crosslinked samples were adjusted to 150 mM NaCl, 0.1 % Tween-20, 10 mM Tris-Cl pH 8.0, 1 mM EDTA and mixed with 10 μl of pre-washed anti-FLAG® M2 Magnetic Beads (Sigma) at 4 °C overnight. Unbound material (supernatant) was removed by pipetting and beads were washed twice with 200 μl 150 mM NaCl, 0.1 % Tween-20, 10 mM Tris-Cl pH 8.0, 1 mM EDTA pH 8.0, once with 200 μl 700 mM NaCl, 0.1 % Tween-20, 10 mM Tris-Cl pH 8.0, 1 mM EDTA pH 8.0, and four times with 100 μl 10 mM Tris-Cl pH 8.0, 1 mM EDTA pH 8.0. Chemical cleavage mapping reactions (100 μl) were carried out directly on beads using the piperidine base-hydrolysis method (Bednar *et al*., 2017). Samples were resuspended in 10 μl formamide, incubated at 90°C for 2 min, then loaded on a prerun acrylamide (6 %)/urea (8 M) sequencing gels and run at 55 W for 65 minutes. Gels were dried and analyzed by autoradiography, on a Typhoon FLA 9500 (GE healthcare).

## References

Allan, J., Mitchell, T., Harborne, N., Bohm, L., and Crane-Robinson, C. (1986). Roles of H1 domains in determining higher order chromatin structure and H1 location. J. Mol. Biol. 187, 591–601.

Bates, D.L., Butler, J.G., Pearson, E.C., and Thomas, J.O. (1981). Stability of the higher-order structure of chicken erythrocyte chromatin in solution. Eur. J. Biochem. 119, 469–476.

Bednar, J., Garcia-Saez, I., Boopathi, R., Cutter, A.R., Papai, G., Reymer, A., Syed, S.H., Lone, I.N., Tonchev, O., Crucifix, C., et al. (2017). Structure and Dynamics of a 197 bp Nucleosome in Complex with Linker Histone H1. Mol Cell 66, 384–397 e388. 10.1016/j.molcel.2017.04.012.

Bondos, S.E., Dunker, A.K., and Uversky, V.N. (2022). Intrinsically disordered proteins play diverse roles in cell signaling. Cell Commun Signal 20, 20. 10.1186/s12964-022-00821-7.

Borgia, A., Borgia, M.B., Bugge, K., Kissling, V.M., Heidarsson, P.O., Fernandes, C.B., Sottini, A., Soranno, A., Buholzer, K.J., Nettels, D., et al. (2018). Extreme disorder in an ultrahigh-affinity protein complex. Nature 555, 61–66. 10.1038/nature25762.

Bradbury, E.M., Cary, P.D., Chapman, G.E., Crane-Robinson, C., Danby, S.E., Rattle, H.W., Boublik, M., Palau, J., and Aviles, F.J. (1975). Studies on the role and mode of operation of the very-lysine-rich histone H1 (F1) in eukaryote chromatin. The conformation of histone H1. Eur J Biochem 52, 605–613.

Burge, N.L., Thuma, J.L., Hong, Z.Z., Jamison, K.B., Ottesen, J.J., and Poirier, M.G. (2022). H1.0 C Terminal Domain Is Integral for Altering Transcription Factor Binding within Nucleosomes. Biochemistry 61, 625–638. 10.1021/acs.biochem.2c00001.

Caterino, T.L., Fang, H., and Hayes, J.J. (2011). Nucleosome linker DNA contacts and induces specific folding of the intrinsically disordered h1 carboxyl-terminal domain. Mol Cell Biol 31, 2341–2348. MCB.05145-11 [pii] 10.1128/MCB.05145-11.

Cerf, C., Lippens, G., Muyldermans, S., Segers, A., Ramakrishnan, V., Wodak, S.J., Hallenga, K., and Wyns, L. (1993). Homo- and heteronuclear two-dimensional NMR studies of the globular domain of histone H1: sequential assignment and secondary structure. Biochemistry 32, 11345–11351.

Clark, D.J., Hill, C.S., Martin, S.R., and Thomas, J.O. (1988). Alpha-helix in the carboxy-terminal domains of histones H1 and H5. EMBO J. 7, 69–75.

Clark, D.J., and Kimura, T. (1990). Electrostatic mechanism of chromatin folding. J. Mol. Biol. 211, 883–896.

Cutter, A.R., and Hayes, J.J. (2015). A brief review of nucleosome structure. FEBS Lett 589, 2914–2922. 10.1016/j.febslet.2015.05.016.

Fang, H., Clark, D.J., and Hayes, J.J. (2012). DNA and nucleosomes direct distinct folding of a linker histone H1 C-terminal domain. Nucleic Acids Res. 40, 1475–1484. gkr866 [pii] 10.1093/nar/gkr866.

Fang, H., Wei, S., Lee, T.H., and Hayes, J.J. (2016). Chromatin structure-dependent conformations of the H1 CTD. Nucleic Acids Res 44, 9131–9141. 10.1093/nar/gkw586.

Hansen, J.C., Lu, X., Ross, E.D., and Woody, R.W. (2006). Intrinsic protein disorder, amino acid composition, and histone terminal domains. J Biol Chem 281, 1853–1856.

Hao, F., Kale, S., Dimitrov, S., and Hayes, J.J. (2021). Unraveling linker histone interactions in nucleosomes. Curr Opin Struct Biol 71, 87–93. 10.1016/j.sbi.2021.06.001.

Hao, F., Mishra, L.N., Jaya, P., Jones, R., and Hayes, J.J. (2022). Identification and analysis of six phosphorylation sites within the Xenopus laevis H1.0 C-terminal domain indicate distinct effects on nucleosome structure. Molecular & Cellular Proteomics In Press. doi.org/10.1016/j.mcpro.2022.100250.

Hao, F., Murphy, K.J., Kujirai, T., Kamo, N., Kato, J., Koyama, M., Okamato, A., Hayashi, G., Kurumizaka, H., and Hayes, J.J. (2020). Acetylation-modulated communication between the H3 N-terminal tail domain and the intrinsically disordered H1 C-terminal domain. Nucleic Acids Res 48, 11510–11520. 10.1093/nar/gkaa949.

Happel, N., and Doenecke, D. (2009). Histone H1 and its isoforms: contribution to chromatin structure and function. Gene 431, 1–12. S0378-1119(08)00568-4 [pii] 10.1016/j.gene.2008.11.003.

Heidarsson, P.O., Mercadante, D., Sottini, A., Nettels, D., Borgia, M.B., Borgia, A., Kilic, S., Fierz, B., Best, R.B., and Schuler, B. (2022). Release of linker histone from the nucleosome driven by polyelectrolyte competition with a disordered protein. Nat Chem 14, 224–231. 10.1038/s41557-021-00839-3.

Kavi, H., Emelyanov, A.V., Fyodorov, D.V., and Skoultchi, A.I. (2016). Independent Biological and Biochemical Functions for Individual Structural Domains of Drosophila Linker Histone H1. J Biol Chem 291, 15143–15155. 10.1074/jbc.M116.730705.

Lee, J., Crickard, J.B., Reese, J.C., and Lee, T.H. (2019). Single-molecule FRET method to investigate the dynamics of transcription elongation through the nucleosome by RNA polymerase II. Methods 159–160, 51-58. 10.1016/j.ymeth.2019.01.009.

Lu, X., Hamkalo, B., Parseghian, M.H., and Hansen, J.C. (2009). Chromatin condensing functions of the linker histone C-terminal domain are mediated by specific amino acid composition and intrinsic protein disorder. Biochemistry 48, 164–172.

Lu, X., and Hansen, J.C. (2004). Identification of specific functional subdomains within the linker histone H10 C-terminal domain. J Biol Chem 279, 8701–8707.

Lu, X., Wontakal, S.N., Kavi, H., Kim, B.J., Guzzardo, P.M., Emelyanov, A.V., Xu, N., Hannon, G.J., Zavadil, J., Fyodorov, D.V., and Skoultchi, A.I. (2013). Drosophila H1 regulates the genetic activity of heterochromatin by recruitment of Su(var)3-9. Science 340, 78–81. 10.1126/science.1234654.

Murphy, K.J., Cutter, A.R., Fang, H., Postnikov, Y., Bustin, M., and Hayes, J.J. (2017). HMGN1 and 2 Remodel Core And Linker Histone Tail Domains Within Chromatin. Nucleic Acids Res 45, 9917–9930. 10.1093/nar/gkx579.

Pan, C., and Fan, Y. (2016). Role of H1 linker histones in mammalian development and stem cell differentiation. Biochim Biophys Acta 1859, 496–509. 10.1016/j.bbagrm.2015.12.002.

Ramakrishnan, V., Finch, J.T., Graziano, V., Lee, P.L., and Sweet, R.M. (1993). Crystal structure of globular domain of histone H5 and its implications for nucleosome binding. Nature 362, 219–224.

Roque, A., Iloro, I., Ponte, I., Arrondo, J.L., and Suau, P. (2005). DNA-induced secondary structure of the carboxyl-terminal domain of histone H1. J Biol Chem 280, 32141–32147.

Roque, A., Teruel, N., Lopez, R., Ponte, I., and Suau, P. (2012). Contribution of hydrophobic interactions to the folding and fibrillation of histone H1 and its carboxy-terminal domain. J Struct Biol 180, 101–109. 10.1016/j.jsb.2012.07.004.

Sottini, A., Borgia, A., Borgia, M.B., Bugge, K., Nettels, D., Chowdhury, A., Heidarsson, P.O., Zosel, F., Best, R.B., Kragelund, B.B., and Schuler, B. (2020). Polyelectrolyte interactions enable rapid association and dissociation in high-affinity disordered protein complexes. Nature communications 11, 5736. 10.1038/s41467-020-18859-x.

Sridhar, A., Orozco, M., and Collepardo-Guevara, R. (2020). Protein disorder-to-order transition enhances the nucleosome-binding affinity of H1. Nucleic Acids Res 48, 5318–5331. 10.1093/nar/gkaa285.

Syed, S.H., Goutte-Gattat, D., Becker, N., Meyer, S., Shukla, M.S., Hayes, J.J., Everaers, R., Angelov, D., Bednar, J., and Dimitrov, S. (2010). Single-base resolution mapping of H1-nucleosome interactions and 3D organization of the nucleosome. Proc Natl Acad Sci U S A 107, 9620–9625. 1000309107 [pii] 10.1073/pnas.1000309107.

Turner, A.L., Watson, M., Wilkins, O.G., Cato, L., Travers, A., Thomas, J.O., and Stott, K. (2018). Highly disordered histone H1-DNA model complexes and their condensates. Proc Natl Acad Sci U S A 115, 11964–11969. 10.1073/pnas.1805943115.

van Holde, K.E. (1989). Chromatin (Springer Verlag).

Vila, R., Ponte, I., Collado, M., Arrondo, J.L., Jimenez, M.A., Rico, M., and Suau, P. (2001). DNA-induced alpha-helical structure in the NH2-terminal domain of histone H1. J Biol Chem 276, 46429–46435. 10.1074/jbc.M106952200.

Ward, J.J., Sodhi, J.S., McGuffin, L.J., Buxton, B.F., and Jones, D.T. (2004). Prediction and functional analysis of native disorder in proteins from the three kingdoms of life. J Mol Biol 337, 635–645. 10.1016/j.jmb.2004.02.002.

Willcockson, M.A., Healton, S.E., Weiss, C.N., Bartholdy, B.A., Botbol, Y., Mishra, L.N., Sidhwani, D.S., Wilson, T.J., Pinto, H.B., Maron, M.I., et al. (2021). H1 histones control the epigenetic landscape by local chromatin compaction. Nature 589, 293–298. 10.1038/s41586-020-3032-z.

Yang, Z., and Hayes, J.J. (2011). The divalent cations Ca2+ and Mg2+ play specific roles in stabilizing histone-DNA interactions within nucleosomes that are partially redundant with the core histone tail domains. Biochemistry 50, 9973–9981. 10.1021/bi201377x.

Yue, H., Fang, H., Wei, S., Hayes, J.J., and Lee, T.H. (2016). Single-Molecule Studies of the Linker Histone H1 Binding to DNA and the Nucleosome. Biochemistry 55, 2069–2077. 10.1021/acs.biochem.5b01247.

Zhou, B.R., Feng, H., Kale, S., Fox, T., Khant, H., de Val, N., Ghirlando, R., Panchenko, A.R., and Bai, Y. (2021). Distinct Structures and Dynamics of Chromatosomes with Different Human Linker Histone Isoforms. Mol Cell 81, 166–182 e166. 10.1016/j.molcel.2020.10.038.

Zhou, B.R., Jiang, J., Feng, H., Ghirlando, R., Xiao, T.S., and Bai, Y. (2015). Structural Mechanisms of Nucleosome Recognition by Linker Histones. Mol Cell 59, 628–638. 10.1016/j.molcel.2015.06.025.

