## Supplemental Information for "Histone H3 tail modifications regulate structure and dynamics of the H1 C-terminal domain within nucleosomes"

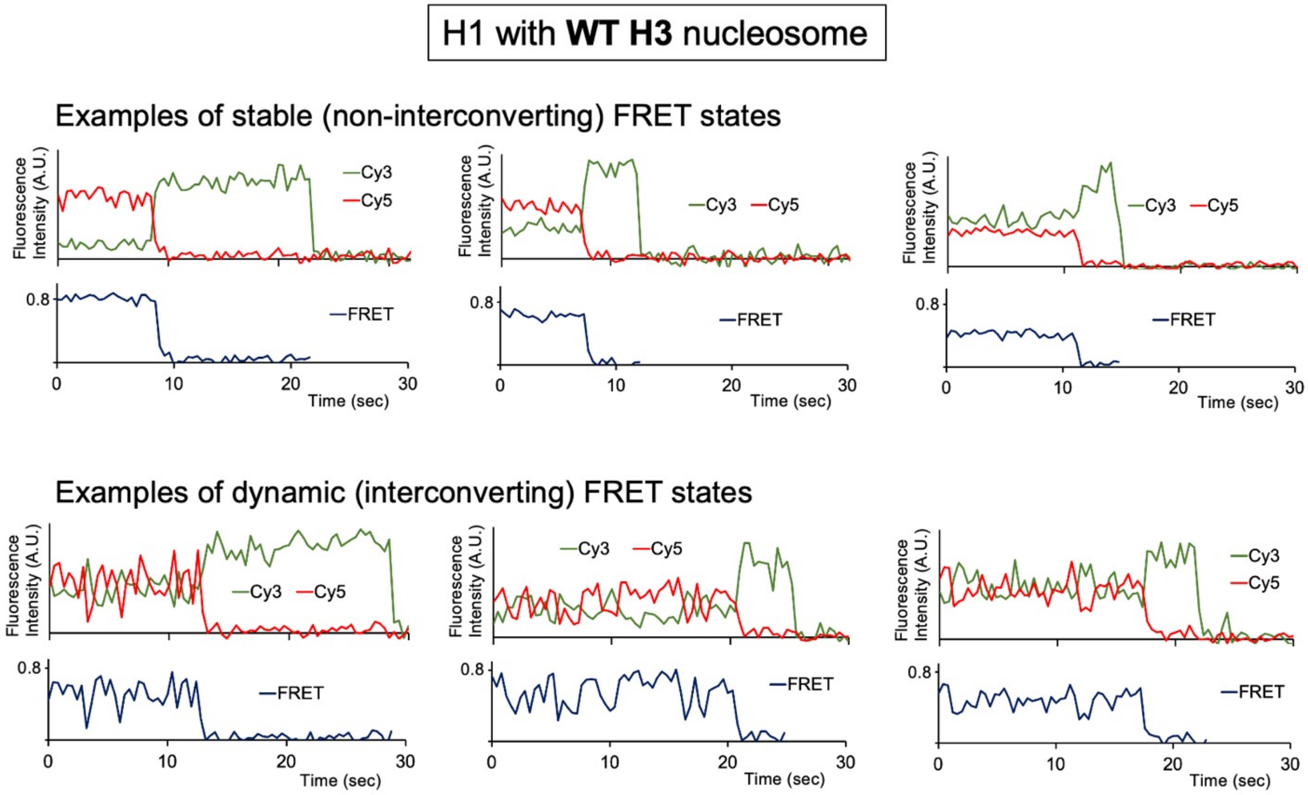


**Fig. S1:** Examples of raw fluorescence intensities and smFRET time traces for static (top) and dynamic (bottom) FRET populations for Cy3/Cy5 labeled H1 G101C K195C bound to WT nucleosomes. Fluorescence intensities are plotted in an arbitrary unit (A.U.).

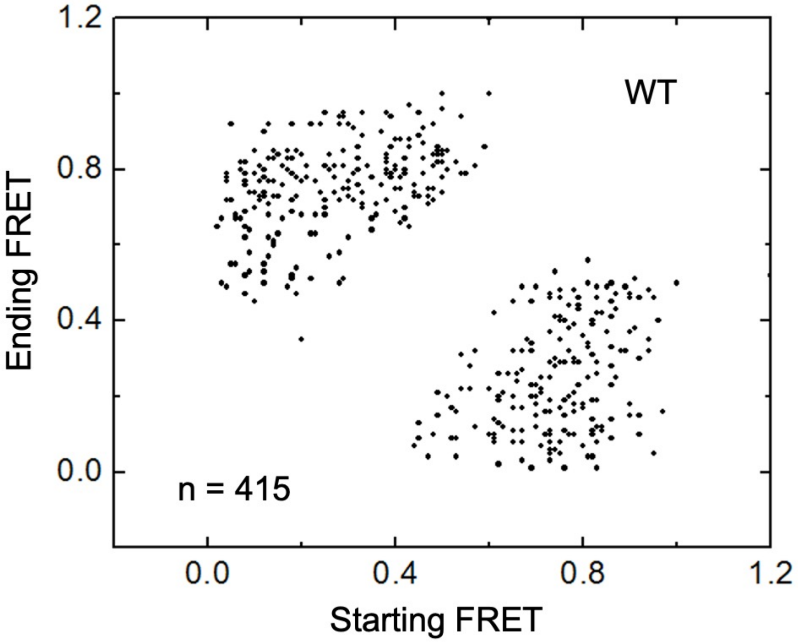


**Fig. S2: FRET transition scatter plot to show no clustering of related conformations.** All FRET time traces used to construct the FRET histograms in WT H1-nucleosome complexes (Fig. 2) were analyzed further to construct a FRET transition scatter plot. The FRET transition time points were identified by visual inspection. The selection criteria are (i) the FRET change should be at least 0.15 and (ii) the intensities of Cy3 and Cy5 are clearly anti-correlated to make matching contributions to the FRET change. The FRET efficiencies immediately before and after each transition point were used to construct this scatter plot. The results clearly show that the transitions do not form any distinct colonies, confirming no clustering of H1 CTD conformations.

a

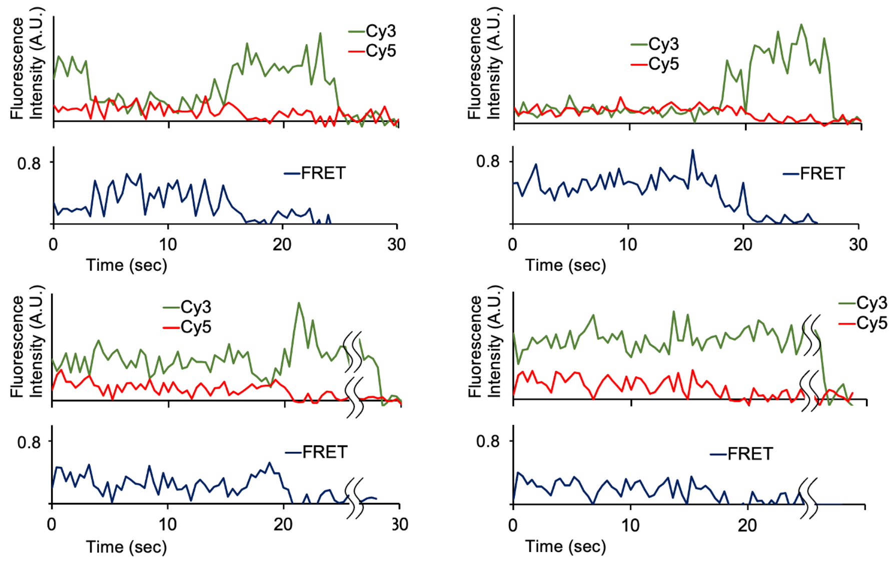


b

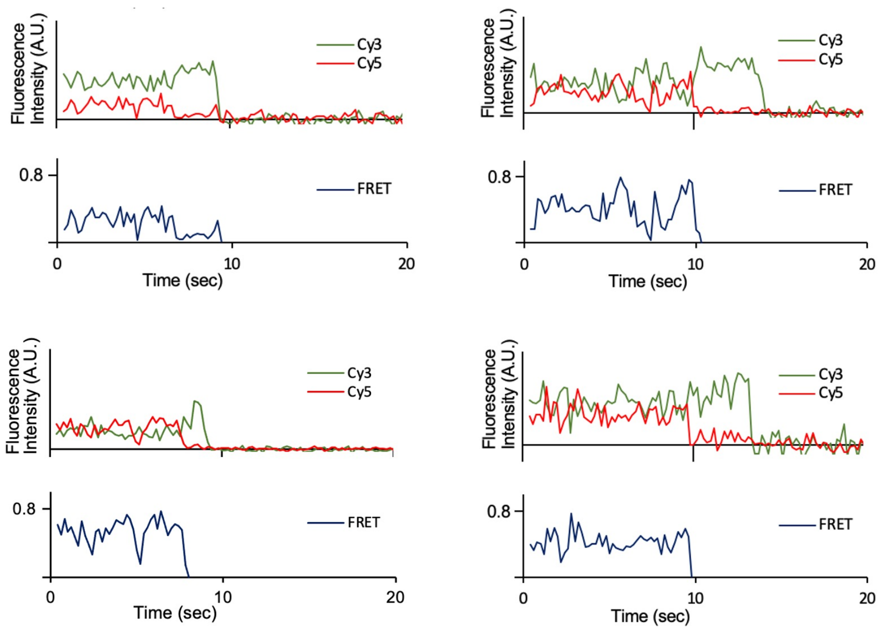


**Fig. S3: Examples of Dynamic FRET traces for free H1s**. Shown are traces for (a) non-specifically bound H1 (see Fig. 1d) and (b) H1 attached via N-terminal FLAG-tag (see text).

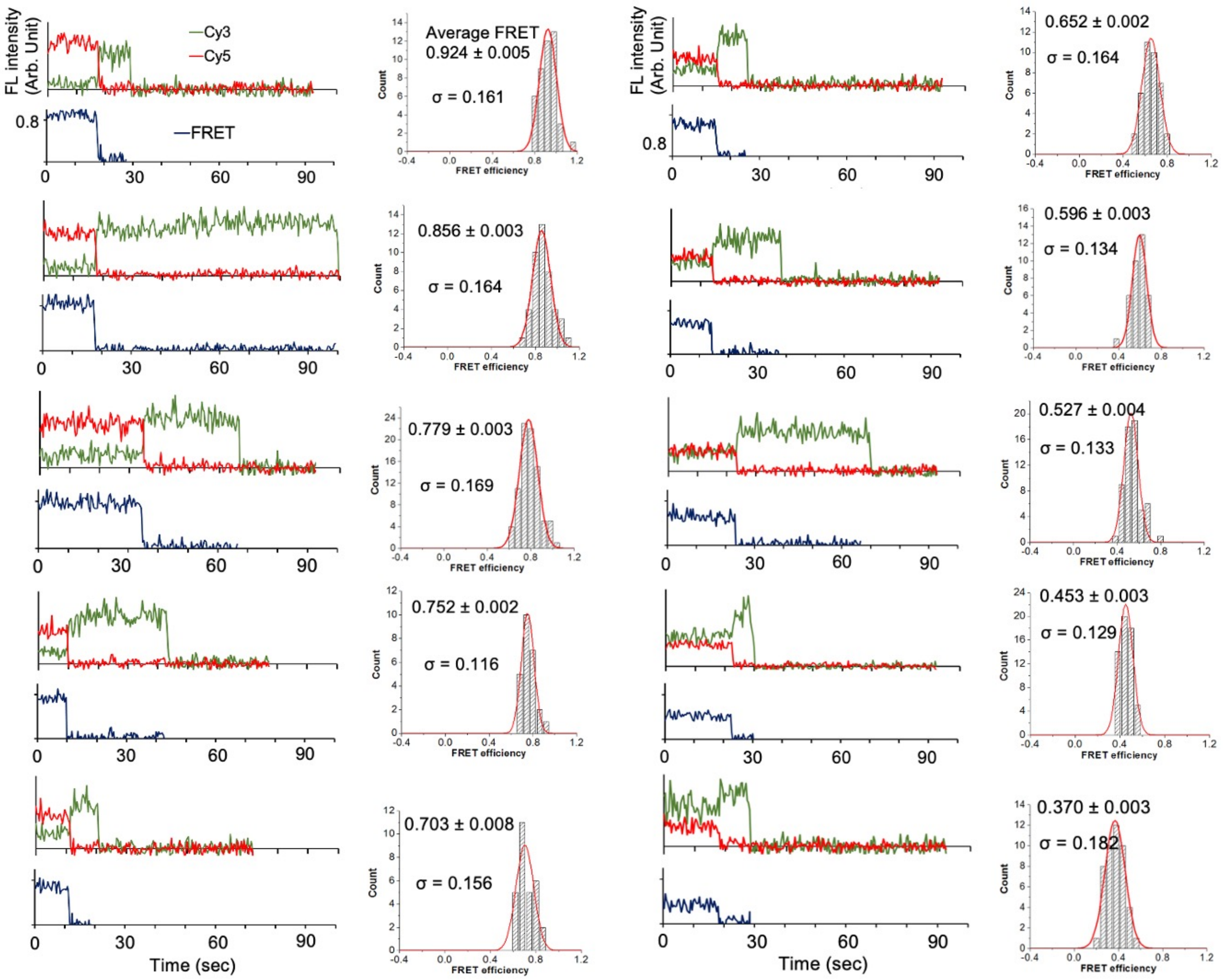


**Fig. S4: Examples of FRET traces show narrow FRET distributions with distinct peak values.** A set of 10 FRET traces were selected from WT H1-nucleosome complexes so that each trace shows a relative Cy3/Cy5 intensity levels distinct from the others. The 10 different relative intensity levels of Cy3/Cy5 in the 10 FRET traces indicate that each trace reports a unique conformation of H1 CTD. The matching FRET histograms are shown below the FRET traces.

WT, N = 162

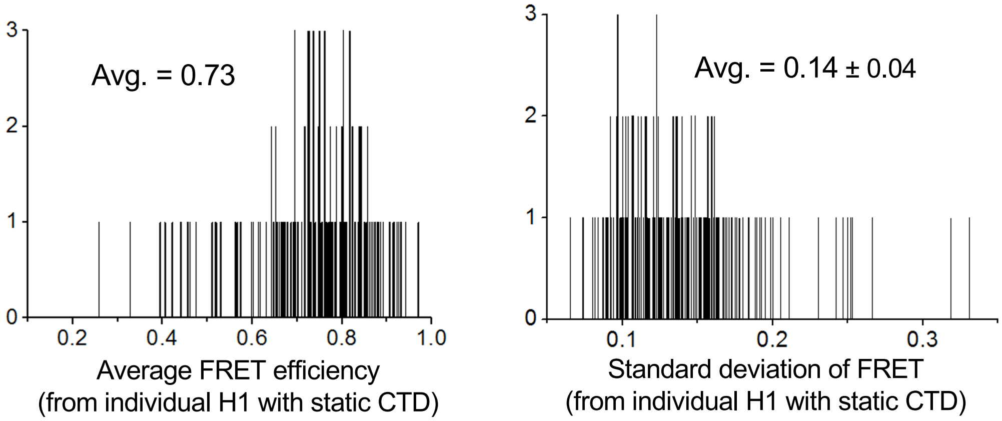


**Fig. S5: Histograms of the average (i.e., peak) and standard deviation of the FRET distributions from the H1 molecules with static CTD associated with WT nucleosomes**

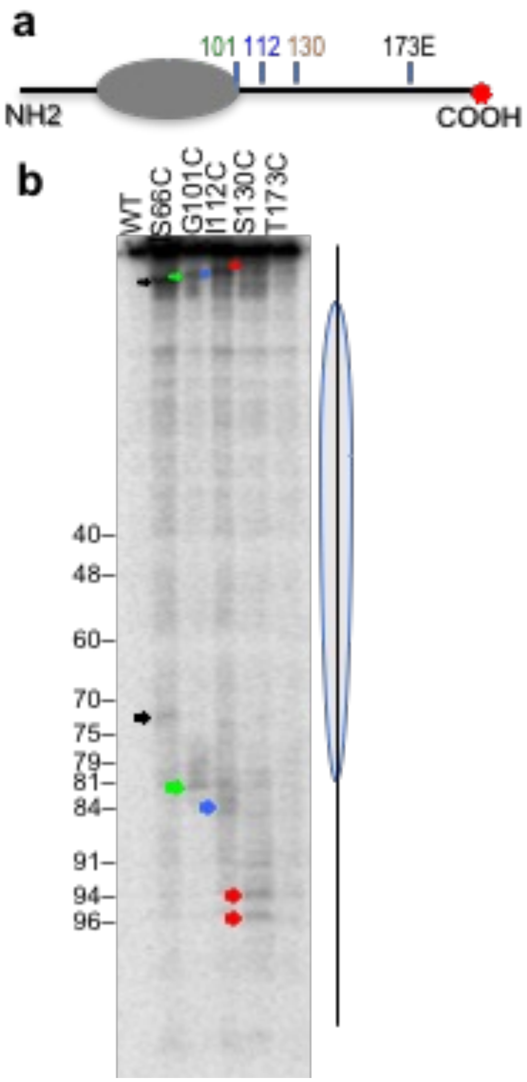


**Fig. S6: Crosslink mapping of sites of association between the H1 CTD and the opposite linker DNA.** a Positions of cysteine substitutions modified with APB in the indicated H1 mutants, as in Fig. 4a. b Crosslink mapping on nucleosome DNA. Sites of crosslinking for S66C-APB. G101C-APB, I112C-APB, S130C-APB are indicated by the black, green, blue and red arrows, respectively.  No discrete crosslinking is detected for T173C-APB. The nucleosome core (oval) and linker DNA (line) are indicated as are positions in the 601 DNA, indicated as distance from the nucleosome dyad position (9). Identical positions are identified to that found for labeling of the 5' end of the top strand (Fig. 4).
